# A plant virus–based virus like-particle platform for therapeutic vaccination targeting IL-23A

**DOI:** 10.64898/2026.01.06.697872

**Authors:** Anete Ogrina-Komarova, Ieva Kalnciema, Dace Skrastina, Arnis Strods, Santa Pikure, Juris Jansons, Gunta Resevica, Janis Bogans, Ramona Petrovska, Andris Zeltins, Ina Balke

**Affiliations:** Plant virology group, Latvian Biomedical Research and Study Centre, Ratsupites street 1, k-1, Riga LV-1067, Latvia; Plant virus protein research group, Latvian Biomedical Research and Study Centre, Ratsupites street 1, k-1, Riga LV-1067, Latvia; Laboratory Animal Core Facility, Latvian Biomedical Research and Study Centre, Ratsupites street 1, k-1, Riga LV-1067, Latvia; Cell Biology and Microscopy Core facility, Latvian Biomedical Research and Study Centre, Ratsupites street 1, k-1, Riga LV-1067, Latvia; Core facility of recombinant protein biotechnology, Latvian Biomedical Research and Study Centre, Ratsupites street 1, k-1, Riga LV-1067, Latvia

**Keywords:** eggplant mosaic virus, virus-like particles, Il-23A, vaccine

## Abstract

Interleukin-23 (IL-23) is a pivotal cytokine in the pathogenesis of immune-mediated inflammatory diseases, making it an attractive therapeutic target. Vaccination against self-antigens offers an alternative to monoclonal antibody (mAb) therapies by eliciting endogenous Ab responses. Here, we report the development of a virus-like particle (VLP)–based vaccine targeting murine IL-23A (mIL23A) using the eggplant mosaic virus (EMV) coat protein as a carrier. mIL23A was incorporated either as a direct fusion or within a mosaic system, with the latter supporting efficient VLP assembly. Alternative antigen presentation strategies employed synthetic coiled-coil and zipper pairs. All constructs were expressed in *E. coli*, purified, and characterized by SDS-PAGE, TEM, and DLS. Immunization of BALB/c mice elicited robust mIL23A-specific Abs, with mosaic VLPs inducing high-avidity, Th1-biased responses. Encapsidated RNA within VLPs was double-stranded and stimulated TLR3, enhancing immunogenicity. Scale-up in bioreactors preserved VLP integrity and antigen incorporation, yielding 1.72 mg per gram of cells. These results establish EMV VLPs as a flexible and scalable platform for presenting challenging antigens and provide a promising approach for therapeutic vaccination against self-antigens.

## Introduction

Innate immunity constitutes the semi-specific first line of defense in both primitive and complex multicellular organisms and provides the initial acute inflammatory response to tissue injury, trauma, or invading pathogens, thereby preventing infection and maintaining homeostasis (Riera Romo et al., 2016). Innate immune mechanisms are capable of mounting an induced response during primary infection by recognizing pathogen-associated molecular patterns (PAMPs) or host-derived damage-associated molecular patterns (DAMPs) through pattern recognition receptors (PRRs). This recognition triggers intracellular signaling cascades leading to the activation of transcription factors such as NF-κB, AP-1, CREB, C/EBP, and IRFs (Newton and Dixit, 2012). Although innate immune responses are rapid and relatively non-specific, they play a critical role in recruiting immune cells to sites of injury or infection via cytokine release, promoting phagocytosis, activating the complement system, and initiating adaptive immune responses (Rubartelli and Lotze, 2007).

Activation of the adaptive immune system results in antigen-specific host responses mediated by T and B lymphocytes (Janeway, 2001). The interplay between innate and adaptive immunity represents a highly coordinated and complex system in which both arms are equally essential for establishing and maintaining immune homeostasis. Each system comprises cellular and humoral components that function synergistically (Nicholson, 2016). Inflammation triggered by pathogens, tissue damage, or trauma is typically self-limiting and resolves with tissue repair following elimination of the inciting cause. However, dysregulation of immune responses may lead to persistent inflammation, progression to chronic inflammatory states, and the development of autoimmune reactions in susceptible individuals. Such altered immune responses are believed to underlie the initiation and progression of immune-mediated inflammatory diseases (IMIDs) (Feghali and Wright, 1997).

IMIDs comprise a heterogeneous group of seemingly unrelated disorders that share common inflammatory pathways. These diseases arise from, or are associated with, dysregulation of innate and adaptive immune system functions and are characterized by disturbances in the physiological cytokine milieu (Kuek et al., 2007, Williams and Meyers, 2002). Although their precise etiology remains unclear, genetic predisposition combined with environmental triggers—such as infections or physical trauma—are thought to initiate disease onset. IMIDs include, but are not limited to, psoriasis, rheumatoid arthritis (RA), inflammatory bowel diseases (IBD), systemic lupus erythematosus, Sjögren syndrome, and type 1 diabetes mellitus (Martin et al., 2014, McGinley et al., 2018). These conditions can affect virtually any organ system and are often associated with reduced quality of life, significant morbidity, and shortened lifespan (Kuek et al., 2007).

Traditionally, Th1 and Th2 subsets of adaptive immune cells were considered the principal drivers of IMID pathogenesis. However, the identification of Th17 cells as a distinct lineage of CD4⁺ helper T cells has substantially advanced our understanding of autoimmunity and inflammation, addressing limitations of the Th1/Th2 paradigm (Bunte and Beikler, 2019). The discovery that interleukin-23 (IL-23) induces the differentiation of naïve precursor cells into Th17 cells further highlighted the importance of this pathway (Harrington et al., 2005). These findings established IL-17 and IL-23 as pivotal cytokines in IMID pathogenesis (Cua et al., 2003).

The pathogenic role of IL-23 was initially demonstrated in experimental autoimmune encephalomyelitis (EAE), a murine model of multiple sclerosis (MS), where IL-23—but not IL-12—was shown to be essential for disease development (Oppmann et al., 2000). This observation was further supported by studies demonstrating that adoptive transfer of IL-17–producing T cells into healthy mice was sufficient to induce EAE (Langrish et al., 2005). Moreover, IL-23–deficient animals failed to recruit IL-17–producing T cells (Murphy et al., 2003), and mice lacking the IL-23–specific p19 subunit (IL-23A), but not the IL-12–specific p35 subunit, were resistant to EAE induction (Cua et al., 2003).

In psoriasis, which was initially regarded as a predominantly Th1-mediated disease, identification of the IL-23/Th17 axis revealed its central role in driving inflammatory progression and provided novel therapeutic targets (Chhabra et al., 2016, Mease, 2015). Beyond dermatological disorders, IL-17 has been implicated as a key mediator in shaping a pathogenic bacterial environment within the oral microbiota (Graves et al., 2019). In a leukocyte adhesion deficiency type I (LAD-I) mouse model of periodontitis, IL-23–dependent IL-17 production promoted bacterial overgrowth, whereas inhibition of this pathway reduced dysbiosis, linking IL-17 overexpression to periodontal disease pathogenesis (Hajishengallis and Moutsopoulos, 2016).

Similarly, elevated IL-23A levels in synovial fluid have been associated with joint destruction in RA through interactions with IL-17, TNF-α, and IL-1β (Kim et al., 2007). Synovial activation in RA is driven by both cytokine-dependent and cytokine-independent mechanisms, including Toll-like receptor (TLR) signaling and endogenous retroviral elements (Ospelt et al., 2004). IL-17 has been shown to upregulate TLR2, TLR4, and TLR9 expression in the synovium of collagen-induced arthritis (CIA) mouse models (Lee et al., 2009), while IL-23 synergizes with IL-17 to enhance TLR expression, underscoring the importance of the IL-23/IL-17 axis in joint inflammation(Lee et al., 2014).

The etiology of IBD, including ulcerative colitis (UC) and Crohn’s disease (CD), remains incompletely understood. Dysbiosis of the intestinal microbiome, combined with genetic susceptibility, is thought to contribute to disease onset by disrupting epithelial barrier integrity and activating aberrant immune responses (Zhang and Li, 2014). The identification of Th17 cells marked a paradigm shift toward recognizing the IL-23/Th17 axis as a central pathogenic pathway in IBDs (Fujino et al., 2003, Yen et al., 2006, Kobayashi et al., 2008, Nielsen et al., 2003). Elevated protein and mRNA levels of IL-23, IL-17, IL-21, and IL-22 have been detected in inflamed intestinal tissues from CD patients, and lamina propria mononuclear cells from these patients secrete increased IL-17 upon T-cell receptor stimulation. Genome-wide association studies further support this pathway, identifying multiple CD-associated polymorphisms in genes related to IL-23/Th17 signaling. Consistently, animal models of intestinal inflammation show reduced disease severity when the IL-23/Th17 pathway is disrupted (Schmitt et al., 2021, Siakavellas and Bamias, 2012).

Spondyloarthropathies (SpA) represent a group of inflammatory arthritides affecting the axial skeleton and/or peripheral joints, with ankylosing spondylitis (AS) as the prototypical form. Extra-articular manifestations, including uveitis, psoriasis, and IBD, are common and form part of SpA classification criteria. Proposed pathogenic mechanisms linking SpA and IBD include gut microbiome dysregulation and migration of immune cells from the intestine to joint tissues (Fragoulis et al., 2019). Notably, entheses contain IL-23 receptor–expressing innate lymphoid cells (ILCs) that express RORγt and CD3 but lack CD4 and CD8 (Sherlock et al., 2012)..In animal models, IL-23 overexpression induces SpA-like phenotypes through activation of these ILCs, which produce IL-17 and IL-22—cytokines implicated in systemic inflammation and pathological bone formation, respectively (Cua and Sherlock, 2011).

Emerging evidence also implicates the IL-23/Th17 axis in other autoimmune and inflammatory disorders, including Hashimoto’s thyroiditis, asthma, obstructive sleep apnea, and allergic airway inflammation (Gerenova et al., 2019, Yi et al., 2022, Lambrecht and Hammad, 2014, Lee and Park, 2022).

IL-23 is a member of the IL-12 cytokine family and is a heterodimer composed of a 19 kDa α subunit (IL-23p19, IL-23A) disulfide-linked to a 40 kDa β subunit (IL-12p40) (Oppmann et al., 2000). The p19 subunit is expressed by antigen-presenting cells, T cells, and endothelial cells, whereas expression of p40 is largely restricted to monocytes, macrophages, and dendritic cells. The formation of biologically active IL-23 requires the co-expression of both subunits within the same cell (Langrish et al., 2005, van de Vosse et al., 2003). IL-12 and IL-23 share the p40 subunit and signal through the IL-12Rβ1 receptor chain expressed on T cells and natural killer (NK) cells (Torti and Feldman, 2007).

Human IL-23p19 shares approximately 70% structural homology with murine p19 and also exhibits homology with the p35 subunit of IL-12. IL-23 signals through a heterodimeric receptor complex consisting of IL-23R and IL-12Rβ1, but not IL-12Rβ2. The IL-23R chain, which directly binds IL-23A, comprises an extracellular N-terminal immunoglobulin-like domain followed by two cytokine receptor domains and belongs to the haematopoietin receptor family (Parham et al., 2002). In contrast, IL-12Rβ1 contains three membrane-proximal fibronectin type III domains and two cytokine receptor domains that interact with the shared p40 subunit (van de Vosse et al., 2003). IL-23R expression is predominantly observed on activated memory T cells and is also detected on NK cells and, at lower levels, on monocytes/macrophages and dendritic cells. IL-12Rβ1, in turn, is broadly expressed on T cells, NK cells, and dendritic cells. In addition to forming functional IL-12 and IL-23 heterodimers, the p40 subunit can also form homodimers. These p40 homodimers have been shown to induce lymphotoxin-α production in microglia and macrophages via IL-12Rβ1, but not IL-12Rβ2 (Jana and Pahan, 2009). Based on these observations, Tang *et al*. proposed that IL-12Rβ1 signaling is dominant in autoimmune inflammation compared with IL-12Rβ2 (Tang et al., 2012). Collectively, these findings suggest that IL-23 may promote autocrine and paracrine signaling loops within the innate immune system, leading to sustained production of inflammatory mediators (McGovern and Powrie, 2007). IL-23 thus serves as a critical cytokine bridging innate and adaptive immunity (Langrish et al., 2004) and plays an essential role in driving early local immune responses (McKenzie et al., 2006).

IL-23 was initially reported to induce IFN-γ production, a cytokine central to Th1 responses and cell-mediated immunity against intracellular pathogens (Yen et al., 2006, Langrish et al., 2005, Oppmann et al., 2000). More recent studies have expanded the functional repertoire of IL-23, demonstrating its involvement in tumor growth and metastasis. IL-23 can directly bind IL-23R expressed on cancer cells in several inflammation-associated malignancies, including oral, lung, liver, and colorectal cancers (Fukuda et al., 2010, Li et al., 2012, Li et al., 2013, Zhang et al., 2014).

The treatment of immune-mediated inflammatory diseases (IMIDs) has traditionally relied on glucocorticoids, non-steroidal anti-inflammatory drugs, and disease-modifying antirheumatic drugs such as methotrexate. Although these therapies are effective in alleviating clinical symptoms and remain first-line treatments, issues related to intolerance, insufficient efficacy, and severe adverse effects have driven the development of alternative therapeutic strategies. Consequently, monoclonal antibodies (mAbs) and fusion proteins—collectively referred to as biologics—were introduced in the early 1990s (Bunte and Beikler, 2019). Clinical trials demonstrated improved disease outcomes following treatment with anti-p40 Abs (Kauffman et al., 2004, Mannon et al., 2004).

Therapies targeting IL-17 or the upstream cytokine IL-23 have shown remarkable efficacy, particularly in psoriasis. Abs targeting IL-17A, IL-17RA, IL-23p19, and IL-12p40 have been approved for the treatment of moderate-to-severe plaque psoriasis (Teng et al., 2015). Although efficacy in multiple sclerosis has been more limited, promising results have emerged from early clinical trials using the anti-IL-17A Ab secukinumab (Havrdová et al., 2016). IL-23–targeted therapies have also demonstrated efficacy in palmoplantar pustulosis, with guselkumab approved for clinical use in Japan (Terui et al., 2019), and have shown potential in systemic sclerosis (Fukasawa et al., 2022), plaque psoriasis, and psoriatic arthritis (Kerut et al., 2023). Risankizumab, a selective anti-IL-23p19 Ab, is approved for moderate-to-severe plaque psoriasis, active psoriatic arthritis, and CD (Feagan et al., 2017, Kristensen et al., 2022, Pang et al., 2024). Other anti-IL-23 Abs, including mirikizumab and brazikumab, have also been evaluated for CD (Kaneko and Sakuraba, 2025, Sands et al., 2017). Currently, four mAbs—risankizumab, mirikizumab, brazikumab, and guselkumab—are either approved or in advanced clinical trials for CD or UC (Barnes, 2024, Parigi et al., 2022). A fifth mAb, tildrakizumab, targeting IL-23p19, has been approved for plaque psoriasis and evaluated for psoriatic arthritis (Mease et al., 2021, Papp et al., 2015, Reich et al., 2017).

Investigation of anti-IL-23 therapies in other autoimmune diseases continues to expand. Anti-IL-23A treatment significantly reduced IL-17A, IL-6, and TNF-α expression in the aortas of ApoE⁻/⁻ mice (Wang et al., 2019). In murine EAE models, therapeutic blockade of IL-23A decreased serum IL-17 levels, reduced central nervous system expression of IFN-γ, IP-10, IL-17, IL-6, and TNF mRNA, prevented inflammatory cell infiltration, and ultimately inhibited disease development. Notably, anti-IL-23A Abs did not increase infection risk to the same extent as anti-p40 Abs and were associated with reduced cancer risk in murine tumor models (Tang et al., 2012). These findings validate the IL-23–driven IL-17A pathway as a critical therapeutic target in T cell–mediated autoimmunity (McGinley et al., 2018).

Despite their success, biologic therapies are associated with several limitations. A common adverse effect is the development of anti-drug Abs, which can lead to primary or secondary treatment resistance. Furthermore, therapeutic efficacy remains incomplete for many patients; for example, in RA, targeted therapies induce six-month remission in only 25–30% of patients, while partial or non-response is observed in up to 70% within the first six months of treatment. In addition, biologics are costly and require frequent long-term administration (Assier et al., 2017).

In this context, therapeutic vaccination against cytokines has emerged as a promising alternative approach (Semerano et al., 2012). This strategy induces endogenous production of therapeutic Abs, offering the advantages of improved tolerability and the absence of xenogeneic epitopes. A prototype vaccine targeting TNF-α (TNF-K), generated by coupling human TNF-α to keyhole limpet hemocyanin (KLH), demonstrated efficacy in animal models of arthritis and progressed through several clinical trials (Le Buanec et al., 2006). Importantly, vaccinated animals did not develop hypersensitivity to intracellular pathogens such as *Listeria monocytogenes* or *Mycobacterium tuberculosis* (Assier et al., 2016). Vaccination strategies targeting IL-23 have also been explored using IL-23A-derived peptides. Given the potentially opposing roles of IL-12 and IL-23 in autoimmune models, selective inhibition of IL-23 is critical. In collagen-induced arthritis and colitis models, IL-23A peptide vaccines linked to KLH or fused to hepatitis B core antigen demonstrated protective effects (Assier et al., 2017). Plant virus coat protein–derived virus-like particles (VLPs) represent a promising carrier platform for IL-23A–based vaccines and have proven to be versatile tools in nanobiotechnology and vaccine development (Balke et al., 2025, Balke and Zeltins, 2019).

Our studies and others have demonstrated the utility of plant VLPs for vaccines targeting self-antigens, allergens, respiratory infectious diseases, intracellular parasites, and cancer, inducing robust and neutralizing Ab responses (Bachmann et al., 2018, Zeltins et al., 2017), allergens (Josi et al., 2024, Krenger et al., 2024b, Mohsen et al., 2022, Mohsen et al., 2021, Sobczak et al., 2023, Zeltins et al., 2017). Among these platforms, eggplant mosaic virus (EMV) has emerged as a novel and effective carrier for antigen presentation (Ogrina et al., 2023). EMV is a 30 nm icosahedral virus with *T=3* symmetry belonging to the genus *Tymovirus* (Martelli et al., 2002). Its positive-sense single-stranded RNA genome encodes three open reading frames, with ORF3 encoding the 19.9 kDa coat protein (CP), which is translated from a subgenomic RNA (Osorio-Keese et al., 1989).

Using the major cat allergen Fel d 1 as a model antigen, we previously demonstrated that EMV CP can support VLP assembly through multiple antigen presentation strategies, including direct genetic fusion, mosaic expression, chemical coupling, and co-expression using synthetic zipper pairs (SYNZIP 18/17) or coiled-coil–forming peptides (Ecoil/Kcoil). All approaches preserved VLP integrity and elicited high titers of Fel d 1–specific Abs in immunized mice (Ogrina et al., 2023).

In the present study, we developed four murine IL-23A (mIL-23A)–containing EMV-based vaccine variants using mosaic expression, SYNZIP 18/17, SYNZIP 1/2, and Ecoil/Kcoil systems. All constructs formed VLPs, albeit with varying levels of antigen incorporation, enabling evaluation of low and intermediate antigen densities on self-antigen immunogenicity. Direct fusion of mIL-23A to EMV CP disrupted VLP assembly despite partial solubility of the fusion protein. Immunization studies revealed that lower antigen incorporation did not limit immunogenicity but rather influenced optimal vaccination regimens. Mosaic EMV (mEMV)–mIL-23A induced high-avidity Abs with potency to neutralize mIL-23, accompanied by Ab class switching, potentially driven by encapsidated host RNA (Krenger et al., 2024a, Sobczak et al., 2024). Notably, mIL-23A could not be expressed in *E. coli* as a standalone protein or as fusions with SZ2, SZ17, or Kcoil; however, fusion to EMV CP and co-expression with AZ1, SZ18 or Ecoil modified CP markedly enhanced solubility. This effect parallels that of classical solubility-enhancing fusion tags such as maltose-binding protein (MBP) and glutathione S-transferase (GST) commonly used in bacterial protein expression systems (Maina et al., 1988, Smith and Johnson, 1988).

## Results

### mIL23A containing EMV CP VLP construction and characterization

To generate EMV CP–derived VLPs displaying murine IL-23A (mIL-23A) as a target antigen, we used a previously validated EMV CP construct containing a C-terminal flexible 15-amino-acid glycine–serine linker [(GGGGS)_3_; G4S] (Ogrina et al., 2023). As an initial approach, a direct fusion construct comprising EMV-CG4S and the mature mIL-23A peptide (EMV-CG4S-mIL-23A) was generated and expressed in *Escherichia coli* strain C2566. SDS–PAGE analysis revealed weak expression of the expected ∼40.1 kDa fusion protein (Fig 1C). Following test purification by sucrose density gradient centrifugation (Fig 1 A,B,C), sucrose cushions (Fig 1D), transmission electron microscopy (TEM) and dynamic light scattering (DLS) analyses demonstrated that the direct fusion protein formed aggregates rather than VLPs (Fig 1E,F).

**Figure 1.**
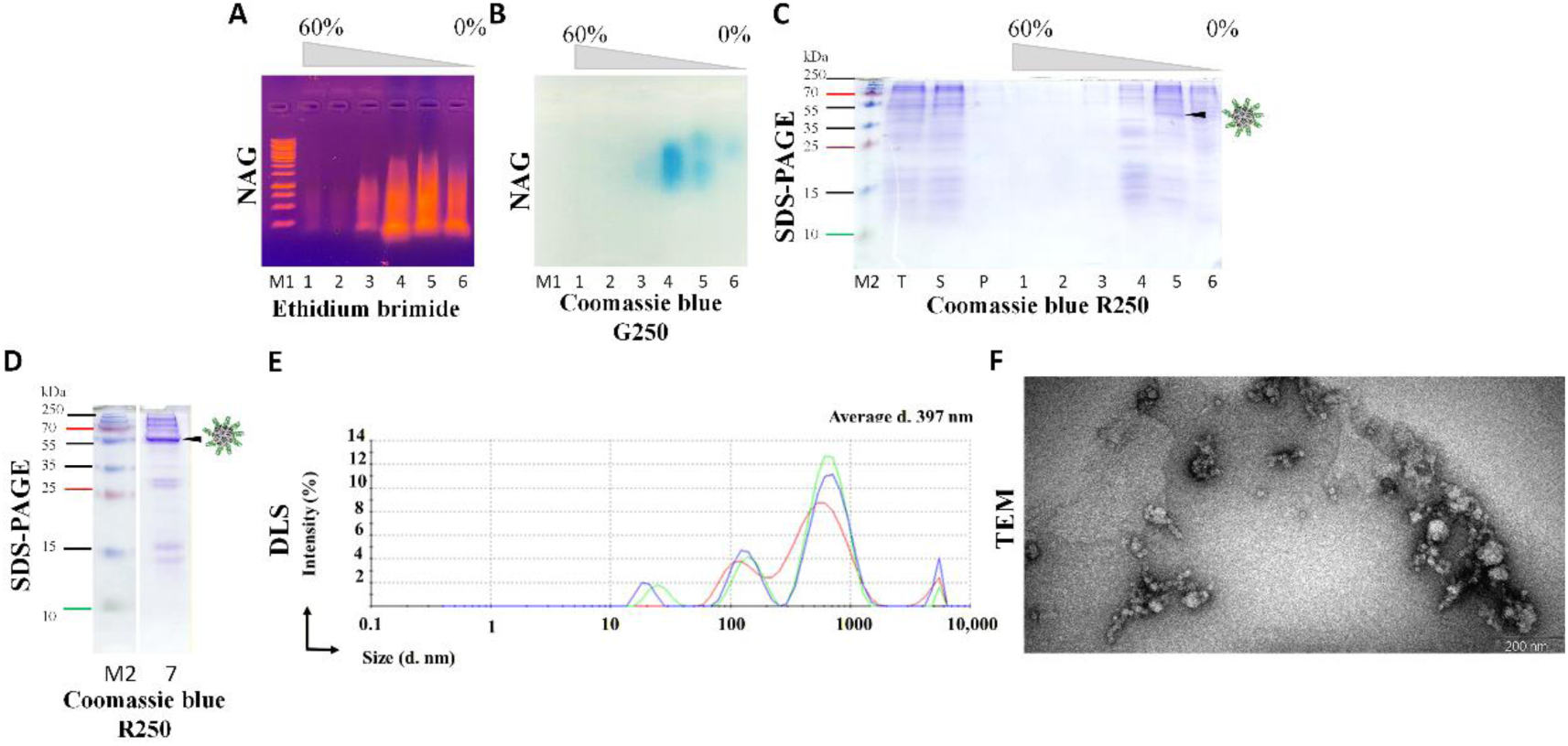
Expression, purification, and characterization of EMV-CG4S-mIL23A VLPs. (**A**) Native agarose gel electrophoresis (NAG) analysis of sucrose density gradient fractions stained with ethidium bromide: T, total cell lysate after 16 h expression; S, soluble fraction after cell disruption and clarification; P, insoluble fraction; lanes 1–6, sucrose gradient fractions starting from 60% sucrose; M1, GeneRuler DNA Ladder Mix (Thermo Fisher Scientific, #SM033).(**B**) NAG analysis of sucrose density gradient fractions stained with Coomassie blue G250 (lane designations as in B). (**C**) SDS-PAGE analysis of sucrose density gradient fractions stained with Coomassie blue R250 (lane designations as in B); M2, PageRuler™ Plus Prestained Protein Ladder (10–250 kDa). (**D**) Analysis of purified EMV-CG4S-mIL23A using a 12.5% SDS-PAGE stained with Coomassie blue R250. (**E**) Dynamic light scattering (DLS) analysis of purified EMV-CG4S-mIL23A VLPs. (**F**) Transmission electron microscopy (TEM) of purified EMV-CG4S-mIL23A VLPs negatively stained with 1% aqueous uranyl acetate. Scale bar: 200 nm. Arrows indicate band corresponding to the expressed fusion protein.

Based on our previous experience with plant VLP-based vaccines targeting peanut allergen Ara h 2 (Sobczak et al., 2023) and cat allergen Fel d 1 (Ogrina et al., 2023, Ogrina et al., 2022), we next employed a mosaic expression strategy. This approach involved co-expression of unmodified wild-type (WT) EMV CP and the EMV-CG4S-mIL-23A fusion protein from a single pETDuet-1 vector (mEMV-CG4S-mIL-23A). Expression in *E. coli* yielded two distinct protein bands corresponding to WT EMV CP (∼19.8 kDa) and EMV-CG4S-mIL-23A (∼40.9 kDa), as confirmed by SDS–PAGE (Fig 2A,E).

**Figure 2.**
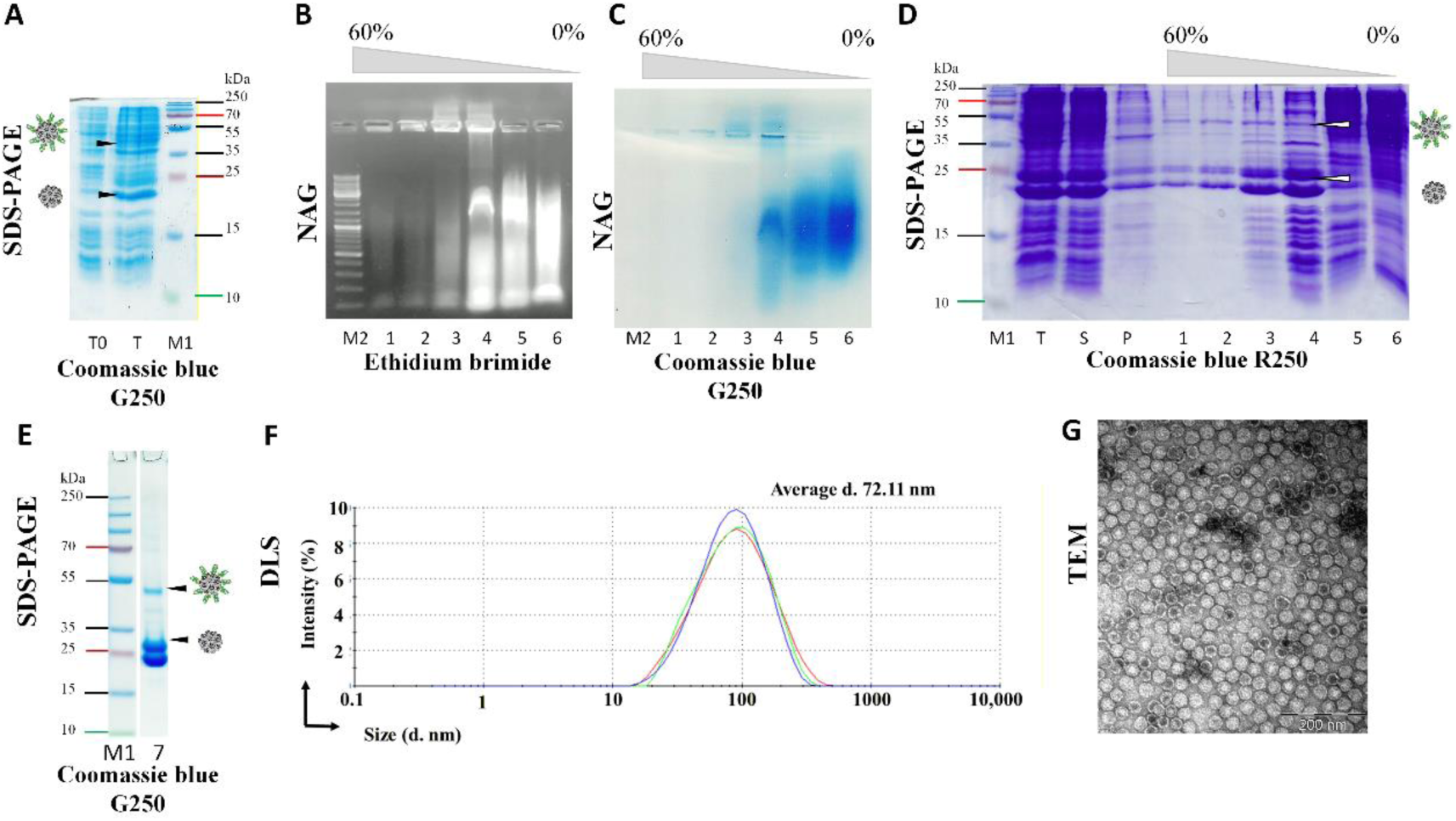
Expression, purification, and characterization of mEMV-CG4S-mIL23A VLPs. (**A**) SDS-PAGE analysis of mEMV-CG4S-mIL23A expression after 16 h: T0, total cell lysate before induction; T, total cell lysate after 16 h of expression; M1, PageRuler™ Plus Prestained Protein Ladder (10–250 kDa; Thermo Fisher Scientific, #26620). (**B**) Native agarose gel electrophoresis (NAG) analysis of sucrose density gradient fractions stained with ethidium bromide: T, total cell lysate after 16 h expression; S, soluble fraction after cell disruption and clarification; P, insoluble fraction; lanes 1–6, sucrose gradient fractions starting from 60% sucrose; M2, GeneRuler DNA Ladder Mix (Thermo Fisher Scientific, #SM033). (**C**) NAG analysis of sucrose density gradient fractions stained with Coomassie blue G250 (lane designations as in B). (**D**) 12.5% SDS-PAGE analysis of sucrose density gradient fractions stained with Coomassie blue R250 (lane designations as in B); M, PageRuler™ Plus Prestained Protein Ladder (10–250 kDa). (**E**) Analysis of purified mEMV-CG4S-mIL23A using a 1.0 mm 4–12% Bis-Tris gel stained with Coomassie blue G250. (**F**) Dynamic light scattering (DLS) analysis of purified mEMV-CG4S-mIL23A VLPs. (**G**) Transmission electron microscopy (TEM) of purified mEMV-CG4S-mIL23A VLPs negatively stained with 1% aqueous uranyl acetate. Scale bar: 200 nm. Arrows indicate bands corresponding to the expressed fusion protein.

Purified mEMV-CG4S-mIL-23A by sucrose gradient (Fig 2 B,C,D) and two sucrose cushions (Fig E) preparations were analyzed by TEM and DLS, revealing uniform *Tymovirus*-like particles with an average diameter of ∼30 nm (Fig 2F,G). Densitometric analysis of SDS–PAGE gels indicated an approximate mIL-23A incorporation rate of 22% within mosaic VLPs. The yield after purification was approximately 22 mg per liter of flask culture (2.55 mg per gram of cells).

As alternative antigen display strategies, we employed three binding-partner systems based on coiled-coil interactions: the Kcoil/Ecoil pair derived from SNARE proteins (Kumar et al., 2018, Litowski and Hodges, 2002), and two orthogonal antiparallel synthetic zipper pairs, SYNZIP17/18 and SYNZIP1/2 (Reinke et al., 2010, Thompson et al., 2012). The selected binding partners form stable physical complexes between protein partners when Kcoil/Ecoil, SZ17/18 or SZ1/2 sequences are genetically introduced in target proteins. Previously generated constructs encoding EMV-CG4S-Ecoil and EMV-CG4S-SZ18 (Ogrina et al., 2023) were used, and a new EMV-CG4S-SZ1 construct was generated for this study.

Correspondingly, mIL-23A was cloned into expression vectors encoding the complementary positively charged partners Kcoil, SZ17, or SZ2. The resulting binding-partner expression vectors were co-expressed in *E. coli* strain C2566. Co-expression variants were purified using the same procedures as applied for the direct and mosaic fusion constructs.

The EMV-CG4S-SZ18 (26.2 kDa) and SZ17-mIL-23A (26 kDa) protein pair exhibited very similar molecular weights, which limited the discrimination of individual components by SDS–PAGE (Fig. 3). However, the conjugated complex could be detected in purified samples as a weak band with an apparent molecular mass of approximately 52.2 kDa (Fig 3E).

**Figure 3.**
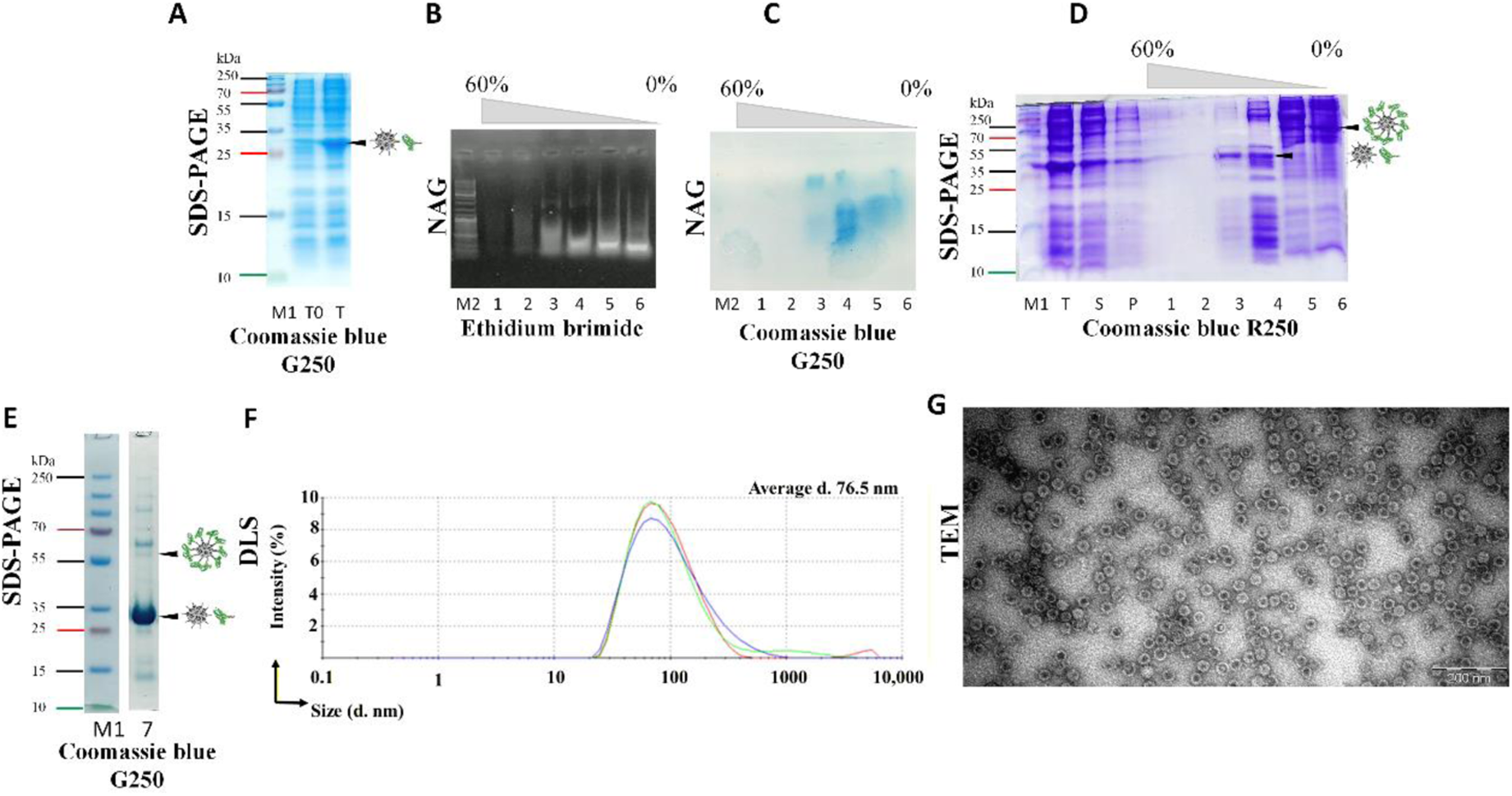
Expression, purification, and characterization of EMV-CG4S-SZ18/SZ17-mIL-23A VLPs. (**A**) SDS-PAGE analysis of EMV-CG4S-SZ18/SZ17-mIL-23A expression after 16 h: T0, total cell lysate before induction; T, total cell lysate after 16 h of expression; M1, PageRuler™ Plus Prestained Protein Ladder (10–250 kDa; Thermo Fisher Scientific, #26620). (**B**) Native agarose gel electrophoresis (NAG) analysis of sucrose density gradient fractions stained with ethidium bromide: T, total cell lysate after 16 h expression; S, soluble fraction after cell disruption and clarification; P, insoluble fraction; lanes 1–6, sucrose gradient fractions starting from 60% sucrose; M2, GeneRuler DNA Ladder Mix (Thermo Fisher Scientific, #SM033). (**C**) NAG analysis of sucrose density gradient fractions stained with Coomassie blue G250 (lane designations as in B). (**D**) 12.5% SDS-PAGE analysis of sucrose density gradient fractions stained with Coomassie blue R250 (lane designations as in B); M, PageRuler™ Plus Prestained Protein Ladder (10–250 kDa). (**E**) Analysis of purified mEMV-CG4S-mIL23A using a 1.0 mm 4–12% Bis-Tris gel stained with Coomassie blue G250. (**F**) Dynamic light scattering (DLS) analysis of purified EMV-CG4S-SZ18/SZ17-mIL-23A VLPs. (**G**) Transmission electron microscopy (TEM) of purified EMV-CG4S-SZ18/SZ17-mIL-23A VLPs negatively stained with 1% aqueous uranyl acetate. Scale bar: 200 nm. Arrows indicate bands corresponding to the expressed fusion protein.

Similarly, EMV-CG4S-SZ1 (25.4 kDa) with SZ2-mIL-23A (26.8 kDa), and EMV-CG4S-Ecoil (23.5 kDa) with Kcoil-mIL-23A (22.7 kDa), also displayed closely overlapping molecular weights, resulting in merged bands in SDS–PAGE (Fig 4A,D,E; Fig 5A,D,E). In these cases, weak higher-molecular-weight bands corresponding to the conjugated complexes (∼52.2 kDa and ∼46.2 kDa, respectively) were also detectable (Fig. 4E; Fig 5E). TEM and DLS analyses demonstrated the formation of VLPs with sizes comparable to those observed for mEMV-CG4S-mIL-23A (Fig 3F,G; Fig 4F,G; Fig 5F,G). Densitometric analysis of SDS–PAGE images indicated antigen incorporation ratios of 0.3% for EMV-CG4S-SZ1/SZ2-mIL-23A, 1.56% for EMV-CG4S-SZ18/SZ17-mIL-23A, and 3.4% for EMV-CG4S-Ecoil/Kcoil-mIL-23A. The estimated yields were approximately 30 mg per liter of flask culture (3.44 mg per gram of cells) for EMV-CG4S-SZ1/SZ2-mIL-23A, 29.8 mg per liter (3.62 mg per gram of cells) for EMV-CG4S-SZ17/SZ18-mIL-23A, and 19.8 mg per liter (1.75 mg per gram of cells) for EMV-CG4S-Ecoil/Kcoil-mIL-23A.

**Figure 4.**
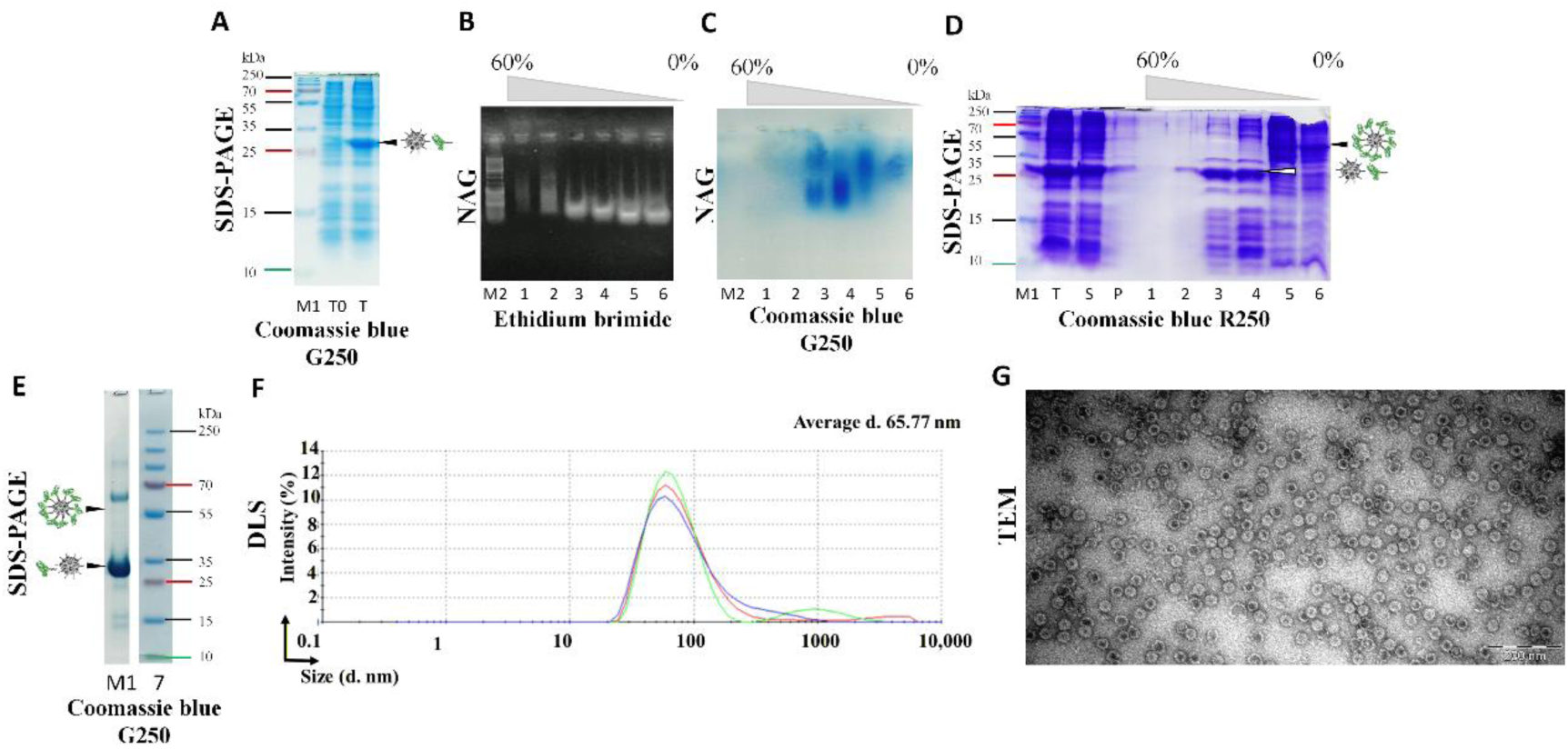
Expression, purification, and characterization of EMV-CG4S-SZ1/SZ2-mIL-23A VLPs. (**A**) SDS-PAGE analysis of EMV-CG4S-SZ1/SZ2-mIL-23A expression after 16 h: T0, total cell lysate before induction; T, total cell lysate after 16 h of expression; M1, PageRuler™ Plus Prestained Protein Ladder (10–250 kDa; Thermo Fisher Scientific, #26620). (**B**) Native agarose gel electrophoresis (NAG) analysis of sucrose density gradient fractions stained with ethidium bromide: T, total cell lysate after 16 h expression; S, soluble fraction after cell disruption and clarification; P, insoluble fraction; lanes 1–6, sucrose gradient fractions starting from 60% sucrose; M2, GeneRuler DNA Ladder Mix (Thermo Fisher Scientific, #SM033). (**C**) NAG analysis of sucrose density gradient fractions stained with Coomassie blue G250 (lane designations as in B). (**D**) 12.5% SDS-PAGE analysis of sucrose density gradient fractions stained with Coomassie blue R250 (lane designations as in B); M, PageRuler™ Plus Prestained Protein Ladder (10–250 kDa). (**E**) Analysis of purified mEMV-CG4S-mIL23A using a 1.0 mm 4–12% Bis-Tris gel stained with Coomassie blue G250. (**F**) Dynamic light scattering (DLS) analysis of purified EMV-CG4S-SZ1/SZ2-mIL-23A VLPs. (**G**) Transmission electron microscopy (TEM) of purified EMV-CG4S-SZ1/SZ2-mIL-23A VLPs negatively stained with 1% aqueous uranyl acetate. Scale bar: 200 nm. Arrows indicate bands corresponding to the expressed fusion protein.

**Figure 5.**
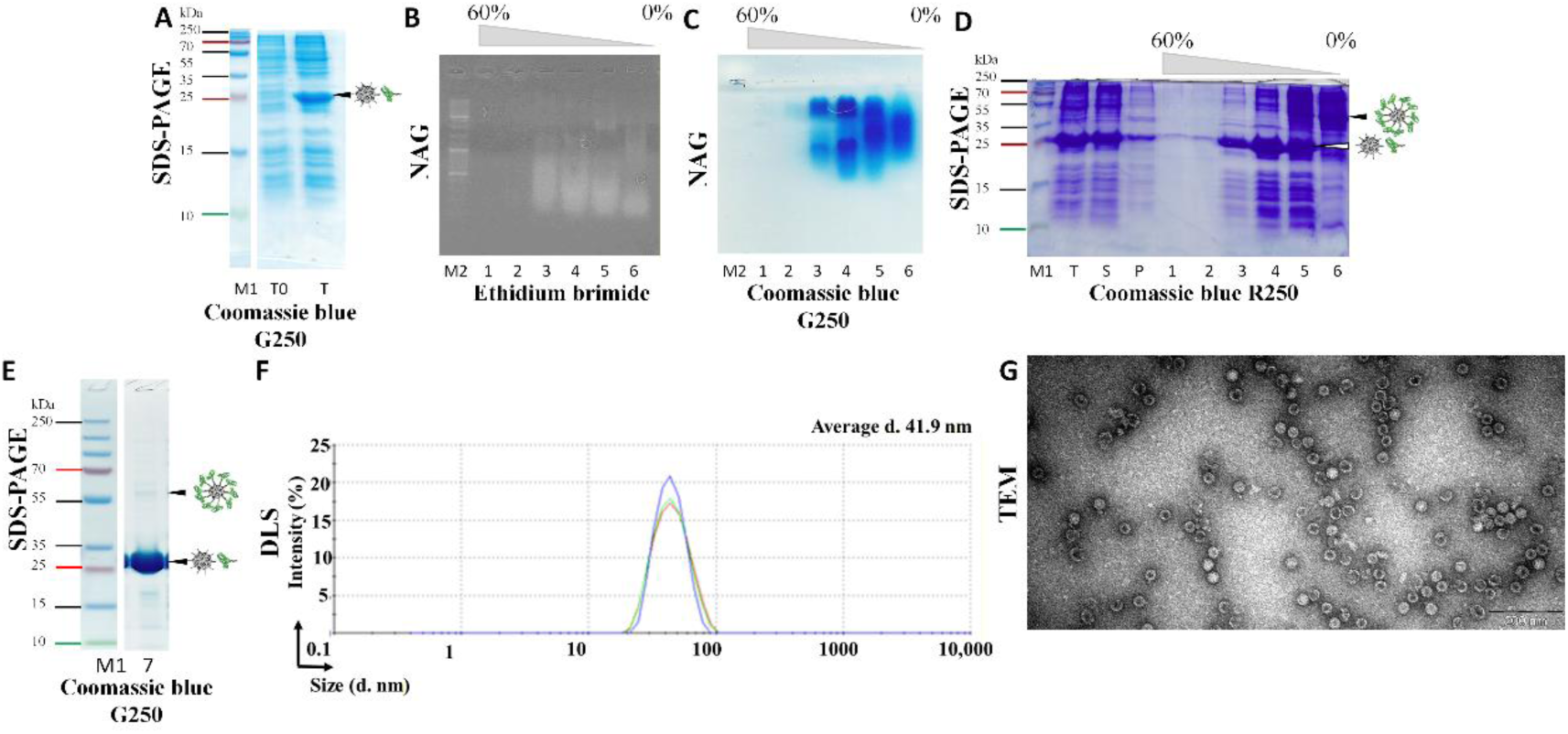
Expression, purification, and characterization of EMV-CG4S-Ecoil/Kcoil-mIL-23A VLPs. (**A**) SDS-PAGE analysis of EMV-CG4S-Ecoil/Kcoil-mIL-23A expression after 16 h: T0, total cell lysate before induction; T, total cell lysate after 16 h of expression; M1, PageRuler™ Plus Prestained Protein Ladder (10–250 kDa; Thermo Fisher Scientific, #26620). (**B**) Native agarose gel electrophoresis (NAG) analysis of sucrose density gradient fractions stained with ethidium bromide: T, total cell lysate after 16 h expression; S, soluble fraction after cell disruption and clarification; P, insoluble fraction; lanes 1–6, sucrose gradient fractions starting from 60% sucrose; M2, GeneRuler DNA Ladder Mix (Thermo Fisher Scientific, #SM033). (**C**) NAG analysis of sucrose density gradient fractions stained with Coomassie blue G250 (lane designations as in B). (**D**) 12.5% SDS-PAGE analysis of sucrose density gradient fractions stained with Coomassie blue R250 (lane designations as in B); M, PageRuler™ Plus Prestained Protein Ladder (10–250 kDa). (**E**) Analysis of purified mEMV-CG4S-mIL23A using a 1.0 mm 4–12% Bis-Tris gel stained with Coomassie blue G250. (**F**) Dynamic light scattering (DLS) analysis of purified EMV-CG4S-Ecoil/Kcoil-mIL-23A VLPs. (**G**) Transmission electron microscopy (TEM) of purified EMV-CG4S-Ecoil/Kcoil-mIL-23A VLPs negatively stained with 1% aqueous uranyl acetate. Scale bar: 200 nm. Arrows indicate bands corresponding to the expressed fusion protein.

### Immunological analysis of mIL23A targeting vaccine prototypes

The immunogenicity of the mIL-23A vaccine candidates was evaluated in female BALB/c mice (7–8 weeks old). Animals were divided into four groups (*n* = 6 per group) and immunized subcutaneously (s.c.) with 30 μg of either mEMV-CG4S-mIL-23A, EMV-CG4S-SZ1/SZ2-mIL-23A, EMV-CG4S-SZ18/SZ17-mIL-23A, or EMV-CG4S-Ecoil/Kcoil-mIL-23A at 14-day intervals (Fig 6A). Serum samples were collected prior to immunization (D0), after the first boost (D14), after the second boost (D28), and at the experimental endpoint (D42).

**Figure 6.**
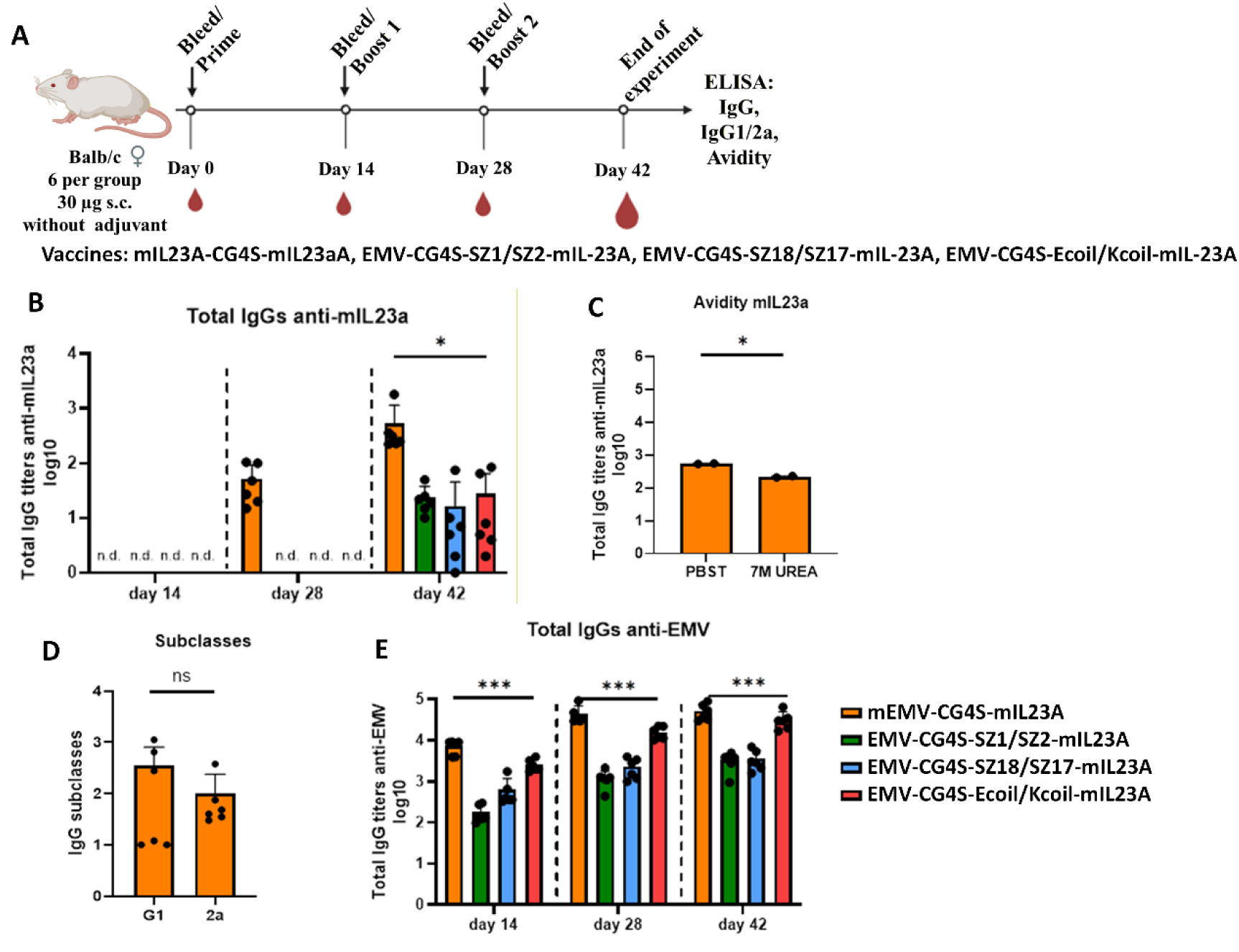
mIL23A vaccine canditate imunological analysis by ELISA. (A) Vaccination regimen and bleeding schedule. (B) Log10 values (mean ± SEM, *n* = 6) of anti-mIL23A specific IgG titers for the groups vaccinated with EMV-mIL23A VLPs on D14, D28, and D4. ELISA plates were coated with 10 µg of CMV-mIL23A per well. (C) Log10 values (mean ± SEM, *n* = 6) of anti-EMV-specific IgG titers for the groups vaccinated with EMV-mIL23A VLPs on D14, D28, and D42. ELISA plates were coated with 10 µg of WT EMV per well. (D) Log10 values of mIL23A specific IgG1 and IgG2 a titers measured in D42 mice sera; ELISA plates were coated with 0.5 µg of recombinant mouse IL-23 protein (Active; amino acids 22–196; Abcam, #ab259423); (E) Avidity antibody detection after vaccination with mEMV-CG4S-mIL23A vaccine Log10 values (mean ± SEM, *n* = 2) of mIL23A specific IgG titers. ELISA plates were coated with 10 µg of carrier EMV CP per well. Statistical analys is using Student’s *t*-test in GraphPad Prism 10.6.1. Vaccine groups *n* = 6. Graphs are showing the meanvalues of single titers. A value of *p* > 0.05 was considered statistically significant, *p* > 0.05 (*); *p* < 0.0001 (***), ns = not significant.

Total serum IgG Abs specific for mIL-23A or the EMV carrier were quantified by ELISA at Days 14, 28, and 42 (Fig 6B). No detectable mIL-23A-specific Abs were observed in any group following the primary immunization (D14; Fig. 6B). Anti-mIL-23A Abs were detected after the first booster dose (D28) exclusively in mice immunized with mEMV-CG4S-mIL-23A, whereas mice receiving EMV-CG4S-SZ1/SZ2-mIL-23A, EMV-CG4S-SZ18/SZ17-mIL-23A, or EMV-CG4S-Ecoil/Kcoil-mIL-23A developed detectable Ab responses only after the second booster at the end of the experiment (D42; Fig 6B).

Avidity assays were performed solely for the mEMV-CG4S-mIL-23A group. Sera from all six animals were pooled and analyzed in duplicate. High-avidity Abs were assessed by washing ELISA plates with 7 M urea to remove low-avidity interactions. Approximately 42% of the mIL-23A-specific Abs induced by mEMV-CG4S-mIL-23A exhibited moderate avidity (Fig 6C).

To further characterize the immune response of mEMV-CG4S-mIL-23A, IgG1 and IgG2a subclass titers were measured in sera collected on D42. A clear dominance of IgG1 over IgG2a was observed among mIL-23A-specific Abs, indicating a predominantly IgG1-biased response (Fig 6D).

Abs directed against the EMV VLP carrier were also detected in all immunized groups beginning at D14, albeit at varying levels (Fig 6E). Tag-based VLP variants elicited lower carrier-specific IgG responses than the mosaic VLP construct. No EMV-specific Abs were detected prior to immunization. Despite high carrier-specific Ab titers in the mosaic group, avidity indices against EMV were low, suggesting preserved particle structure and surface properties. Collectively, these findings indicate that mIL-23A-specific IgG responses are strongly influenced by antigen abundance and spatial organization on the VLP surface. Encapsidated VLP-associated RNA appears to significantly enhance immunogenicity, compensating for reduced antigen repetitiveness in low-incorporation variants and restoring effective immune activation (Krenger et al., 2024a). The residual immunogenicity of the EMV carrier further indicates maintenance of overall VLP integrity and surface charge.

### EMV carrier RNA and mEMV-CG4S-mIL23A VLPs stimulate TLR3

The absence of TLR7 signaling can be partially compensated by TLR3-mediated activation, as both receptors are expressed in B cells (Browne, 2012; Marshall-Clarke et al., 2007). TLR3 is a pattern recognition receptor that senses double-stranded RNA (dsRNA) and may therefore contribute to the observed immunostimulatory effects (Pasare and Medzhitov, 2005). Here, we investigated whether the EMV carrier encapsidates dsRNA and whether intact VLPs are capable of activating TLR3 signaling.

To address this, HEK-Blue™ TLR3 reporter cells were stimulated with 250 or 500 ng of RNA isolated from mEMV-CG4S-mIL23A or WT EMV VLPs using TRI REAGENT, as well as with intact mEMV-CG4S-mIL23A or WT EMV VLPs. Total RNA analysis in NAGE revealed that the main RNA size is under 100 nt (Fig 7A). The total RNA yield differed between the two VLP preparations: RNA isolated from WT EMV VLPs reached 906 ng/µL, whereas RNA obtained from mEMV-CG4S-mIL23A VLPs was approximately 2.26-fold lower (400 ng/µL). Despite this difference, RNA isolated from both VLP types induced dose-dependent activation of TLR3 reporter cells (Fig 7B). Notably, WT EMV-derived RNA elicited significantly stronger TLR3 activation than RNA isolated from mEMV-CG4S-mIL23A VLPs (Fig 7B).

**Figure 7.**
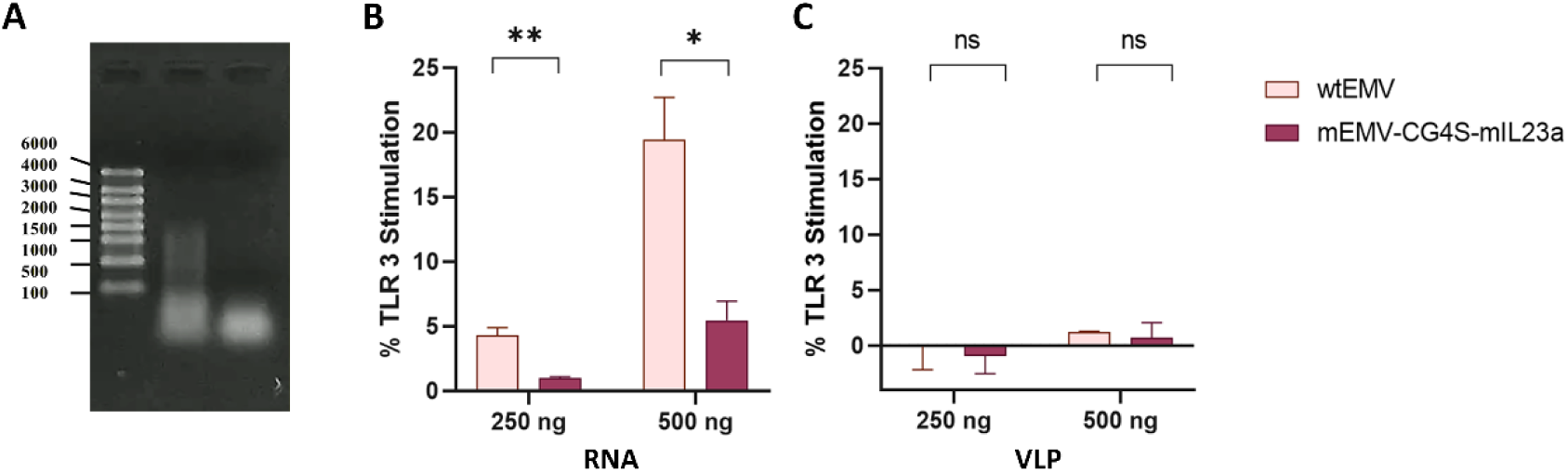
EMV VLP–associated RNA and intact VLP stimulation of TLR3. (**A**) Agarose gel electrophoresis of RNA isolated from VLPs: lane 1, RNA derived from WT EMV VLPs (400 ng/µL); lane 2, RNA derived from mEMV-CG4S-mIL23A VLPs (400 ng/µL); M, RiboRuler High Range RNA Ladder (Thermo Fisher Scientific, #SM1823). (**B**) TLR3 activation in HEK-Blue™ TLR3 reporter cells following stimulation with RNA isolated from WT EMV or mEMV-CG4S-mIL23A VLPs using TRI Reagent. (**C**) TLR3 activation following stimulation with intact WT EMV or mEMV-CG4S-mIL23A VLPs. Results are normalized to positive control stimulated HEK-Blue™ TLR3 cells. Statistical analysis (mean ± SEM) using unpaired Student’s t-test. Ns – not significant, *p* < 0.5 (*), *p* < (**), *n* = 3.

Stimulation of TLR3 reporter cells with intact VLPs also showed a dose-dependent activation pattern (Fig 7C). Higher levels of TLR3 activation may be achieved by increasing the VLP concentration. These results indicate that bacterial RNA encapsidated within both mEMV-CG4S-mIL23A and WT EMV VLPs is present, at least in part, in a double-stranded form capable of triggering TLR3 signaling. Furthermore, intact VLPs are sufficient to activate TLR3 reporter cells, although optimization of VLP concentration may be required. Together, these findings demonstrate that VLPs can be directly used to assess TLR3 ligand activity and suggest a mechanism by which VLP-associated RNA contributes to immune stimulation.

### Scale-up production of mEMV-CG4S-mIL23A

We next assessed the scalability of mEMV-CG4S-mIL23A production toward large-scale vaccine manufacturing. Preliminary results confirmed efficient incorporation of mIL-23A into VLPs, which correlated with a robust antigen-specific immune response. Avidity assays further suggested that the elicited Abs may possess neutralizing activity.

Prior to scale-up of the vaccine candidate, expression of WT EMV CP was tested in one module of a four-module laboratory bioreactor system (EDF 1.1). Final biomass densities (OD_590_) reached 160. SDS-PAGE analysis of total cell lysates before and after induction showed expression patterns comparable to those obtained in flask cultures (Fig 8).

**Figure 8.**
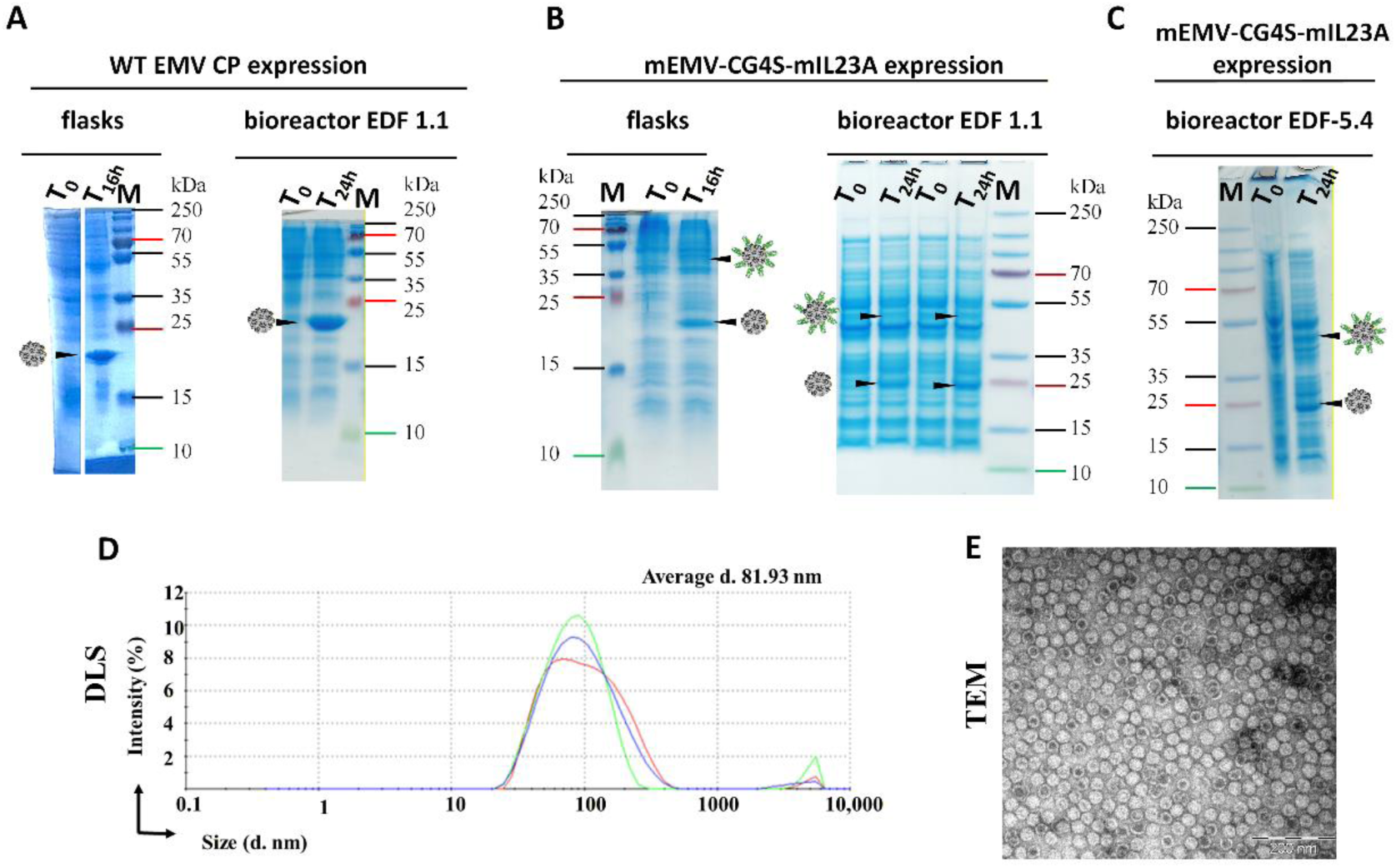
WT EMV CP and mEMV-CG4S-mIL23A scale-up bioreactor. (A) WT EMV CP (19.8 kDa) expression comparison between flask and bioreactor expression: T_0_, total cel lysate befor expression induction; T_16h_, total cell lysat after 16 h expression in flasks; T_24h_, total cell lysate after 24 h expression in bioreactor EDF 1.1; (B) mEMV-CG4S-mIL23A (19.8 kDa, 40.9 kDa) expression comparison between flask and EDF 1.1 bioreactor expression: T_0_, total cel lysate befor expression induction; T_16h_, total cell lysat after 16 h expression in flasks; T_24h_, total cell lysate after 24 h expression in bioreactor EDF 1.1; (C) mEMV-CG4S-mIL23A expression analysi in EDF-5.4 bioreactor: T_0_, total cel lysate befor expression induction; T_24h_, total cell lysate after 24 h expression in bioreactor EDF-5.4. Balck arows indicate corresponding band of expressed protein.

Subsequently, scaled-up production of mEMV-CG4S-mIL23A was evaluated in two 1-L modules of the EDF 1.1 bioreactor. Samples collected before induction (T0) and 24 h post-induction (T24) were analyzed by SDS-PAGE. Final biomass densities (OD_590_) reached 140 and 132 for modules 1 and 2, respectively, and protein expression profiles were consistent with those observed in flask cultures (Fig 8). Further scale-up was performed in a 5-L benchtop fermenter (EDF 5.4). SDS-PAGE analysis of samples collected at T0 and T24 revealed expression patterns comparable to those obtained in both flask and 1-L bioreactor cultures. The final biomass density reached an OD_590_ of 144 (Fig 8).

Cells harvested from the 5-L bioreactor were resuspended in disintegration buffer and processed using a microfluidizer. VLPs were purified using the same protocol as for flask-derived material. SDS-PAGE analysis demonstrated antigen incorporation levels comparable to those observed in VLPs produced in flasks. TEM and DLS analyses confirmed similar particle morphology and size distribution across production scales (Fig 8). The final VLP yield from bioreactor cultures was 1.72 mg per gram of cells.

## Discussion

Early studies identified IL-12 as a key mediator of inflammatory bowel disease (IBD) and other immune-mediated inflammatory diseases (IMIDs); however, the anti-p40 Abs used in these studies neutralized both IL-12 and IL-23 (Yen et al., 2006). The subsequent discovery of Th17 cells and their associated cytokines, IL-17 and IL-23, clarified their central role in autoimmunity and inflammation, highlighting IL-23 as a critical link between innate and adaptive immunity (Bunte and Beikler, 2019, Langrish et al., 2004, McGovern and Powrie, 2007). Conventional IMID therapies, including glucocorticoids, NSAIDs, and methotrexate, are effective but limited by adverse effects and variable efficacy. The introduction of biologics, particularly mAbs targeting IL-17A, IL-17RA, IL-23A, and IL-12p40, has substantially improved outcomes in diseases such as psoriasis (Teng et al., 2015). In CD, however, blockade of downstream IL-17 pathways can exacerbate disease, whereas selective targeting of IL-23A appears safer, with reduced infection and tumor risk observed in preclinical models (Hueber et al., 2012, Tang et al., 2012, Targan et al., 2016). Together, these findings establish the IL-23–IL-17 axis as a validated therapeutic target in IMIDs (McGinley et al., 2018). Vaccination against self-antigens such as IL-23 represents a promising alternative to passive Ab therapy, offering sustained Ab production without xenogenic components. Previous preclinical studies targeting IL-23A or downstream mediators have demonstrated protective effects in IMID models (Assier et al., 2017, Guan et al., 2013, Ratsimandresy et al., 2011, Zeltins et al., 2017). In this study, we demonstrate that the full-length mature mIL-23A peptide (19.7 kDa) can serve as an effective vaccine antigen when presented on plant virus-derived VLPs.

mIL-23A was successfully incorporated into EMV CP either as a direct fusion or using a mosaic expression strategy. While soluble expression was achieved in both cases, VLP assembly occurred only with the mosaic system. This contrasts with a previously reported EMV-based Fel d 1 vaccine, in which both direct fusion and mosaic approaches supported VLP formation (Ogrina et al., 2023). These findings indicate that antigen physicochemical properties, rather than size alone, critically influence CP self-assembly. Although mIL-23A and Fel d 1 are similar in size (19.7 vs. 18.9 kDa), they differ in isoelectric point (IP) and solubility, with mIL-23A (IP 5.85) being poorly soluble and not expressible in *E. coli* without fusion to EMV CP. Increased IP is known to negatively affect protein solubility and expression, independent of codon optimization (Mehlin et al., 2006). Importantly, EMV CP acted as an effective fusion partner, enabling soluble expression of mIL-23A in a bacterial system. This property also facilitated the development of tag-based antigen presentation systems using synthetic zipper pairs (SZ1/2, SZ17/18) and coiled-coil tags (Ecoil/Kcoil). Although antigen incorporation levels were lower, these approaches provide flexible alternatives when direct genetic or chemical fusion is not feasible. Tag-based co-expression and mosaic assembly enable *in vivo* decoration of VLP surfaces during bacterial cultivation through high-affinity interactions (Kd < 10^-8^), minimizing disruption of VLP self-assembly (Reinke et al., 2010, Thompson et al., 2012). The tag-based co-expression and mosaic strategies enable the production of VLPs decorated with the antigen of interest within a single bacterial cell. These approaches facilitate antigen presentation while minimizing the impact of the expressed antigen on VLP spatial organization and self-assembly, thereby providing versatile antigen presentation platforms. All four constructed vaccine systems were successfully expressed in E. coli and purified as antigen-containing VLPs. The presence of both EMV CP VLPs and the mIL-23A antigen in all vaccine variants was confirmed by SDS-PAGE analysis.

The *E. coli* expression system offers a simple, rapid, robust, and cost-effective platform, with recombinant proteins comprising up to 50% of total cellular protein (Francis and Page, 2010), enabling efficient expression and scalable manufacturing. Accordingly, the vaccine candidates described here represent promising prototypes for future mIL-23A–targeted therapeutic vaccines. To our knowledge, this study provides the first report of successful mIL-23A expression in a bacterial system using a plant virus CP as a fusion partner. Consistent with our previous findings, EMV CP VLPs tolerate C-terminal modifications while maintaining correct particle morphology (Ogrina et al., 2023). Immunological analyses demonstrated that all vaccine variants induced mIL-23A–specific Ab responses. Among them, the mosaic mEMV-CG4S-mIL23A platform elicited the most robust immunity, characterized by high-avidity Abs and IgG subclass switching toward a Th1-biased response. Importantly, Abs induced by mosaic VLPs recognized commercially available active mIL-23, underscoring their functional relevance. The superior performance of mosaic VLPs likely reflects their reduced sensitivity to antigen size and structural constraints compared with non-mosaic particles (Pumpens et al., 2016).

The low incorporation levels of mIL-23A observed for the synthetic zipper pairs and the coiled-coil system underscore the critical importance of antigen density and spatial distribution on the VLP surface. This observation is consistent with the PASP concept, which posits that optimal B-cell activation occurs when antigens are displayed on nanoparticles with an inter-epitope spacing of approximately 5–10 nm—a spatial arrangement characteristic of pathogen surfaces. In addition to epitope spacing, surface charge can influence immune recognition by promoting the formation of a protein corona composed of adsorbed serum proteins and natural Abs (Blanco et al., 2015, Vogelstein et al., 1982). It has been well established that efficient B-cell activation depends on the spacing between antigens on viral or VLP surfaces, with antigen spacing being inversely proportional to antigen density (Bachmann and Jennings, 2010, Bachmann and Zinkernagel, 1997, Vogelstein et al., 1982). The impact of antigen abundance and spatial arrangement on immunogenicity has been demonstrated in VLP-based peanut allergy vaccine (Krenger et al., 2024a) and in our recently developed “Immune-tag” platform, where suboptimal antigen distribution on the VLP surface resulted in reduced Ab responses and delayed seroconversion (Sobczak et al., 2024). Importantly, VLP-associated RNA can substantially enhance the immunogenicity of sparsely displayed antigens by compensating for reduced epitope repetitiveness and restoring immune activation (Krenger et al., 2024a). EMV-based VLPs assemble in the cytosol of *E. coli* and therefore encapsidate prokaryotic RNA, including host mRNA and *CP* mRNA (Bachmann and Jennings, 2010, Balke et al., 2025, Carboni et al., 2023, Strods et al., 2015), similar to native viruses that preferentially package their own genomes (Balke et al., 2023). This selective encapsidation is driven by intrinsic structural density and recognition features of viral nucleic acids (Gopal et al., 2014).

The reduced RNA content observed in mEMV-CG4S-mIL23A VLPs compared with WT EMV VLPs may reflect alterations in the cellular transcriptome caused by the presence of antigen-encoding RNA. Detailed characterization of encapsidated RNA by next-generation sequencing, as previously performed for sobemovirus CP-derived VLPs (Balke et al., 2025), would allow precise comparison of RNA composition and length distributions between VLP variants.

NAGE analysis indicated that the RNA encapsidated within mEMV-CG4S-mIL23A VLPs was predominantly shorter than 100 nt. This likely contributed to the predominance of the IgG1 subclass observed, as shorter RNA fragments provide limited ssRNA—particularly polyU-rich sequences—necessary for efficient TLR7 activation (Diebold et al., 2006, Zhang et al., 2017). In contrast, VLP-based peanut vaccines containing longer host-derived RNA fragments (∼500 nt) elicited IgG2a-dominated responses (Krenger et al., 2024a). The shorter RNA in mEMV-CG4S-mIL23A likely results in insufficient ssRNA for robust TLR7 stimulation and a relative lack of dsRNA for TLR3 activation (Krenger et al., 2024a, Sakaniwa et al., 2023, Sobczak et al., 2024). Larger RNA fragments with more ssRNA can enhance TLR7 signaling, promoting IgG class switching toward IgG2a (Gomes et al., 2019). Among murine Ab subclasses, IgG2a is considered most effective against viral infections (Markine-Goriaynoff et al., 2000). In VLP-based vaccines, IgG1 and IgG2a often serve as indicators of T-helper cell polarization: IgG1 reflects Th2 responses, while IgG2a reflects Th1-type immunity (Bachmann and Kündig, 2017). Th1 responses are particularly important for modulating allergen-specific IgG and controlling IgE-mediated reactions. The IgG1/IgG2a ratio for mEMV-CG4S-mIL23A was 0.29, indicating a Th1-polarized response (ratios ≤0.5, Th1; ≥2.0, Th2; 0.5–2.0, mixed) (Feltquate et al., 1997, Pertmer et al., 1996). The observation is consistent with literature showing that TLR ligand presence, such as ssRNA naturally encapsidated into VLPs, influences isotype switching (Jegerlehner et al., 2007, Markine-Goriaynoff et al., 2000).

The dsRNA presence in encapsidated RNA was tested by TLR3 reporter cell assay. TLR3 reporter cells were similarly stimulated by RNA isolated from WT EMV or mEMV-CG4S-mIL23A, confirming the presence of double-stranded RNA within the encapsidated content. Intact VLPs also induced modest TLR3 activation, demonstrating that new vaccine prototypes can be assessed for TLR3 pathway stimulation without RNA extraction. This approach provides a practical means to evaluate innate immune activation in vitro before advancing to animal models.

Despite the lower IgG2a skewing, avidity analysis demonstrated that mEMV-CG4S-mIL23A induced specific Abs, although the AI did not reach the >50–60% threshold for high-avidity Abs. This likely reflects the insufficient TLR7 signaling from short host RNA fragments, while TLR3 activation and the spatial organization of the antigen on the VLP surface may partially compensate to support specific Ab development. Avidity, reflecting the multivalent binding potential of Abs to antigens, is influenced not only by binding strength but also by the geometry of antigen presentation, which constrains polyvalent interactions (Klein and Bjorkman, 2010, Vorup-Jensen, 2012). These interactions are constrained by the geometry of ligand presentation, meaning that the topological arrangement of the antigen-bearing surface directly influences recognition (Vorup-Jensen, 2012). Thus, avidity provides not only a measure of binding strength but also a qualitative assessment of Ab–antigen interactions, encompassing both the number and stability of these interactions. Importantly, higher avidity has been correlated with increased vaccine efficacy, as demonstrated for the RTS,S malaria vaccine (Dobaño et al., 2019).

Carrier-specific Abs can be a concern in repeated vaccine administration, as pre-existing anti-carrier Abs could clear VLPs and reduce boosting of responses against the target antigen. However, in the case of mEMV-CG4S-mIL23A, the AI of anti-EMV Abs was low, indicating largely non-specific interactions. Furthermore, long-term studies with a vaccine targeting equine IL-5 showed sustained efficacy despite repeated administration of the same carrier over five consecutive years (Jonsdottir et al., 2020). Interestingly, VLPs decorated with SZ or Ecoil/Kcoil tags elicited lower anti-carrier Ab titers. This may result from reduced exposure of the VLP surface for Ab binding and the highly negatively charged EMV surface, which limits complement activation and TLR7 engagement, thereby reducing immune cell activation and enhancing the immunotherapeutic potential of the vaccine (Lebel et al., 2017).

Together, these findings highlight that the antigen density, spatial organization, and encapsidated RNA content of VLPs are key determinants of immune responses. The mEMV-CG4S-mIL23A mosaic platform balances these factors, eliciting specific Th1-biased Abs with adequate avidity while mitigating potential anti-carrier interference, supporting its promise as a therapeutic vaccine platform for self-antigens such as mIL-23A.

Scale-up of mEMV-CG4S-mIL23A in bioreactors showed that the vaccine prototype can be produced in high-density cultures, making it suitable for large-scale manufacturing. High-density expression did not compromise antigen incorporation or VLP integrity. Similar results were reported for a SARS-CoV-2 vaccine prototype using cucumber mosaic virus (CMV) as a carrier, mCuMVTT-RBM (Mohsen et al., 2022), highlighting the scalability of the production process. This approach could enable the manufacture of millions of doses in a single 1,000 L bacterial fermenter, an important consideration for vaccine accessibility in less affluent regions (Mohsen et al., 2022).

Overall, the EMV platform serves as a flexible “toolbox” for antigen presentation, allowing both surface display and co-expression of otherwise poorly expressed or insoluble proteins. The robust immune responses achieved, combined with successful scale-up, underscore EMV’s potential as a versatile carrier for multipurpose vaccine development.

## Methods

### Cloning, expression and purification of the EMV and mIL-23A fusion construct for direct and mosaic systems

The coding sequence of the mature mIL-23A peptide (aa 22–196, lacking the 1–21 aa signal peptide; Accession: NP_112542.1) was obtained via gene synthesis in the pUC57 plasmid (Biocat GmbH) without codon optimization. Plasmids encoding EMV CP for direct or mosaic antigen display were described previously (Ogrina et al., 2023). For the direct expression system, the *mIL-23A* sequence was cloned at the C-terminus of EMV CP via a flexible (GGGGS)₃ linker (Maeda et al., 1997) between BamHI and XhoI sites in pET42-EMV-CG4S (Ogrina et al., 2023), generating pET42-EMV-CG4S-mIL23A. For the mosaic system, pETDu-EMV (Ogrina et al., 2023) was used to subclone the *EMV-CG4S-mIL23A* coding sequence under a second *T7* promoter using NdeI and XhoI, generating pETDu-mEMV-CG4S-mIL23A. Both vectors were transformed into E. coli C2655 competent cells (New England Biolabs). Expression and purification followed established protocols (Balke et al., 2025, Balke et al., 2018, Ogrina et al., 2023, Zeltins et al., 2017). In brief Cells were cultured in 200 mL 2×TY medium supplemented with ampicillin (100 μg/mL for pETDu-mEMV-CG4S-mIL23A) or kanamycin (25 μg/mL for pET42-EMV-CG4S-mIL23A) at 30 °C with agitation at 200 rpm. Expression was induced at OD_600_ = 0.8–1.0 using 2 mM IPTG, with medium supplemented with 2 mM CaCl_2_ and 5 mM MgCl_2_, followed by 16 h incubation at 20 °C. Cells were harvested by centrifugation (8,228 × g, 5 min, 5 °C) and stored at –80 °C. Cells were resuspended in disintegration buffer (1× PBS, 1 mM DTT, 5 mM β-ME, 5% glycerol, 10% sucrose) and lysed using an ultrasound disintegrator (UP200S, Hielscher Ultrasonics) at 70% intensity for 16 min (0.5 s pulses). Lysates were clarified by centrifugation (15,557 × g, 10 min, 4 °C). The supernatant was further purified by sucrose density gradient ultracentrifugation (20–60% sucrose in disintegration buffer with 0.5% TX-100; five density steps) at 106,559 × g for 6 h at 18 °C using a swing-out rotor (SW-32, Beckman Coulter). Fractions were analyzed by 12.5% SDS-PAGE and stained with Coomassie blue G250. Fractions containing the fusion protein (30–40% sucrose) were pooled, transferred to 26.3-ml polycarbonate tubes, and diluted with gradient buffer. VLPs were pelleted by ultracentrifugation (183,960 × g, 3 h, 4 °C; fixed-angle rotor Type 70Ti, Beckman Coulter) and resuspended overnight at 4 °C in 10 mL Buffer A (1× PBS, 1 mM DTT, 0.5% glycerol). The solubilized material was clarified (15,557 × g, 10 min, 4 °C) and further purified by two 30% sucrose cushions (183,960 × g, 4 h, 4 °C, Type 70Ti). Final VLP pellets were resuspended in Buffer B (1× PBS, 1 mM DTT, 5% glycerol). Purity was assessed by 12.5% SDS-PAGE or 1 mm 4–12% Bis-Tris gels (Bolt, Thermo Fisher Scientific). VLP integrity was confirmed by 0.8% native agarose gel electrophoresis (NAGE), dynamic light scattering (DLS), and transmission electron microscopy (TEM). Antigen incorporation density was quantified by densitometry using GelAnalyzer 19.1 (available at www.gelanalyzer.com). WT EMV VLPs were expressed and purified as described previously (Ogrina et al., 2023).

### Cloning, expression and purification of the SZ1/SZ2, SZ18/SZ17 an Ecoil/Kcoil partner conjugation variant

Plasmid constructions for pETDuet-1 EMV CP-SZ18 and CP-Ecoil have been described previously (Ogrina et al., 2023). Coding sequences for the synthetic zipper (SZ) pairs SZ1 and SZ2 were obtained from published sources (Reinke et al., 2010) and purchased as gene synthesis products in pUC57 (BioCat). The EMV CP-CG4S coding sequence was cloned between NdeI and XhoI sites at the N-terminus of the SZ1 construct in pUC57-SZ1. The resulting EMV CP-CG4S-SZ1 fragment was then subcloned into pETDuet-1 at NdeI and XhoI sites to generate pETDu-EMV-CG4S-SZ1. The *mIL-23A* coding sequence was cloned at the C-terminus of SZ2, SZ17 (pUC57-SZ17 or pUC57-SZ2), or Kcoil (pACYCDu-Kcoil (Ogrina et al., 2023)) constructs between BamHI and XhoI sites. *SZ1-mIL23A* and *SZ17-mIL23A* fragments were subcloned into pET28a(+) (Novagen). For co-expression, EMV CP-tagged plasmids (pETDu-EMV-CG4S-SZ18, pETDu-EMV-CG4S-SZ1, pET42-EMV-Ecoil) were co-transformed with their respective mIL-23A-tagged partner plasmids (pET-SZ17-mIL23A, pET-SZ2-mIL23A, pACYCDuet-Kcoil-mIL23A) into *E. coli* strain C2566 (New England Biolabs). Cultivation, expression, and purification followed the same protocols established for the EMV-CG4S-mIL23A direct and mosaic systems.

### Negative stain transmission electron microscopy

Purified samples, at concentrations not exceeding 1 mg/mL, were adsorbed onto carbon Formvar–coated copper grids and negatively stained with 1% aqueous uranyl acetate. The detailed protocol has been described previously (Balke et al., 2022). Grids were examined using a JEM-1230 TEM (JEOL) at an accelerating voltage of 80–100 kV, with magnifications of ×80,000 or ×100,000.

### Dynamic light scattering

Purified VLP samples (1 mg/mL) were analyzed by DLS as previously described (Balke et al., 2018). Measurements were performed using a Zetasizer Nano ZS instrument (Malvern Instruments Ltd., Worcestershire, UK) with DLS software version 7.11. For each sample, three sequential measurements were recorded to ensure reproducibility.

### Vaccination of naïve mice

#### Mice

Immunization experiments with mEMV-CG4S-mIL23A, Ecoil/Kcoil, SZ18/SZ17, and SZ1/SZ2 VLPs were performed using naïve female BALB/c mice (*n* = 6 per group, 7–8 weeks old) obtained from the Laboratory Animal Centre, University of Tartu, Estonia. All animal procedures were approved by the Animal Protection Ethics Committee of the Latvian Food and Veterinary Service (Permission No. 89) and were conducted in accordance with Directive 2010/63/EU, as adopted into national legislation.

#### Immunization regiment

The immunogenicity of the VLP constructs was evaluated in mice using 30 µg of total protein per dose in 200 µL PBS, administered s.c. without adjuvants. Each mouse received injections in both flanks (100 µL per site) on day 0 (D0), based on dosing from previous studies with non-adjuvanted VLPs (Sobczak et al., 2024). Mice were divided into four groups: (1) mEMV-CG4S-mIL23A, (2) EMV-CG4S-SZ1/SZ2-mIL23A, (3) EMV-CG4S-SZ18/SZ17-mIL23A, and (4) EMV-CG4S-Ecoil/Kcoil-mIL23A. Booster injections of the same dose were administered on days 14 (D14) and 28 (D28), and the study concluded on day 42 (D42). Serum samples were collected at D0, D14, D28, and at final bleeding on D42.

### Enzyme-linked immunosorbent assay

ELISA was performed to determine total IgG, IgG subclasses, and avidity of vaccine-induced Abs, following established protocols (Ogrina et al., 2023, Ogrina et al., 2022, Sobczak et al., 2024). 96-well plates (Greiner Bio-One) were coated with 10 µg/mL of EMV VLPs, CMV-mIL23 fusion, or 0.5 µg/mL recombinant mouse IL-23 (Abcam, #ab259423) in 50 mM sodium carbonate buffer, pH 9.6, and incubated at 4 °C ON. Plates were washed, blocked with 1% BSA, and incubated with serial dilutions of mouse sera (1:50 for mIL-23A, 1:400 for EMV). HRP-conjugated rabbit anti-mouse IgG (Sigma-Aldrich, #A9044-2ML) was used as secondary Ab (1:10,000). The reaction was developed with OPD substrate and stopped with 1.2 N H_2_SO_4_; absorbance was read at 492 nm. Endpoint titers were defined as the highest dilution exceeding three times the negative control absorbance (Classen et al., 1987, Lardeux et al., 2016).

Isotype-specific ELISA was performed to determine IgG1 and IgG2a levels using the mouse monoclonal isotyping reagent ISO2 (Sigma-Aldrich, #ISO2-1KT) and HRP-conjugated anti-goat/sheep IgG (Sigma-Aldrich, #A9452-1VL).

Avidity of mIL-23A-specific IgG was measured using an extended ELISA protocol incorporating chaotropic washes with 7 M urea (de Souza et al., 2004, Mohsen et al., 2021, Ogrina et al., 2022, Rothen et al., 2022, Wiuff et al., 2002). 96-well plates were coated with 10 µg/mL EMV VLPs, CMV-mIL23 fusion, or 0.5 µg/mL recombinant mouse IL-23 (Abcam, #ab259423) in 50 mM sodium carbonate buffer, pH 9.6, and incubated ON at 4 °C. Prediluted sera (1:20) were added, followed by 1:3 serial dilutions. Plates were washed with PBS containing 0.01% Tween, then incubated three times for 5 min with either 7 M urea in PBS-Tween or PBS-Tween alone. After washes, the AI was calculated as the ratio of titers with and without urea treatment (Correa et al., 2021, Nurmi et al., 2021, Olsson et al., 2019).

### mEMV-CG4S-mIL23A scale up by fed-batch fermentation

One-stage pre-seed culture was prepared by inoculating 100 µL of frozen *E. coli* strain C2566 with pETDu-mEMV-CG4S-mIL23A stock culture into 50 mL of 2x YT broth supplemented with 100 μg mL^−1^ of AMP in 300 mL Erlenmeyer flasks. Culture was incubated at 37 °C and 200 rpm for 8 h, with OD_590_ reaching at least 5.

Fermentation was initially conducted in a 1 L stirred-tank bioreactor (EDF-1.2; Bioreactors.net) and subsequently scaled up to a 5 L system (EDF-5.4; Bioreactors.net). The scale-up maintained constant dissolved oxygen (DO) and aeration (VVM), while the feed rate was increased proportionally to the reactor volume. The sterile-filtered semi-defined medium used for the batch phase contained the following per liter: 5 g Yeast Extract, 6 g (NH_4_)_2_SO_4_, 5 g Na_2_HPO_4_, 3 g KH_2_PO_4_, 0.8 g MgSO_4_ x 7H_2_O, 1 mL of Trace metals solution, 10 g glucose and 0.05 g FeCl_3_ x 6H_2_O (5 mL added separately into bioreactor as a sterile 10 g/L solution). Trace metal solution was as follows: CuCl_2_ x 2H_2_O (0.02 g/L), H_3_BO_3_ (0.01 g/L), NaI (0.2 g/L), MnSO_4_ x H_2_O (0.2 g/L), ZnSO4 x 7H2O (0.04 g/L), and Na_2_MoO_4_ x 2H_2_O (0.04 g/L). The feed medium was a sterile-filtered concentrated glucose solution (400 g/L) supplemented with 30 g/L (NH_4_)_2_SO_4_, 3 g/L MgSO_4_ x 7H_2_O and 10 mL/L of Trace metal solution.

The bioreactor was inoculated at an initial OD_590_ of 0.3. Culture conditions were maintained as follows: temperature controlled at 30 °C, pH at 7.0±0.2 by automatic addition of 12.5% NH_4_OH, aeration with compressed air at 1.0 vvm, foaming mitigated using 10% Antifoam 204 (Sigma-Aldrich) and dissolved oxygen (DO) maintained above 30% by adjusting agitation speed (200-1000 rpm). Off-gas composition (O_2_ and CO_2_) was monitored on EDF-5.4 bioreactor using BlueInOne ferm (BlueSens) gas analyzer.

The fermentation process consisted of an initial batch phase followed by a fed-batch phase after depletion of the initial glucose (indicated by a sharp rise in the dissolved oxygen). Initial feeding was at linearly increasing feed rate from 3.3 mL/h to 37 mL/h per liter of culture (corresponding to a feed rate increase from 1.3 g to 15 g of glucose per hour per liter of culture) for duration of 10 h to rapidly increase cell biomass. OD_590_ measurement above 100 confirms successful cell biomass production, allowing to proceed with cell induction.

Before induction, vessel temperature was lowered down to 20 °C, then IPTG added at 1 mM final concentration. Simultaneously, a constant feed rate of 15 mL/h per liter of culture (corresponding to a feed rate of 4 g glucose per hour per liter of culture) was set and maintained throughout the induction phase, keeping rest of the cultivation parameters unchanged. After 22 to 24 h, fermentation was stopped and cells were harvested by centrifugation and pellets stored at −20 °C for further downstream processing.

### Cell disintegration by benchtop homogenizer

Small-scale cell disruption was performed using benchtop homogenizer LM20 Microfluidizer® Processor. Frozen cell pellets were suspended in ice-cold lysis buffer (1x PBS, 1 mM DTT, 5 mM β-ME, 5% glycerol, 10% sucrose) at a ratio 1:3 (w/v), then passed three times through the microfluidizer at 20 000 psi pressure. Additional washing of homogenizer with lysis buffer was performed to collect all sample after cell lysis, increasing cell-to-buffer ratio up to 1:6. Samples were kept on ice throughout homogenization process, as well as homogenizer’s cooling coil was kept in ice bath to minimize protein degradation in lysed sample. Lysis efficiency was evaluated by measuring OD_590_ throughout homogenization process.

### HEK- Blue™ TLR 3 cell assay

RNA encapsulated within mEMV-CG4S-mIL23A or WT EMV VLPs was isolated from 200 µL of 1 mg/mL VLPs using TRI REAGENT (Sigma-Aldrich, #T9424-100ML) following the manufacturer’s protocol. Purified RNA pellets were dissolved in 25 μL of DEPC-treated water (Thermo Fisher Scientific). RNA concentration was measured using a NanoDrop™ 1000 (Thermo Fisher Scientific), and size distribution was assessed by 1% agarose gel electrophoresis in 1× TBE buffer with ethidium bromide staining. RNA ladders (Thermo Scientific, #SM1821) were included. Before loading, RNA samples were mixed 1:1 with 2× RNA loading dye (Thermo Fisher Scientific), incubated at 70 °C for 10 min, and chilled on ice for 3 min. Nucleic acids were visualized by ethidium bromide staining in NAG. The TLR3-stimulatory capacity of RNA isolated from mEMV-CG4S-mIL23A, WT EMV VLPs, or intact VLPs was evaluated using HEK-Blue™ hTLR3 cells (InvivoGen, #hkb-htlr3) following the manufacturer’s instructions. 50,000 cells per well were stimulated in triplicate with 250 or 500 ng of RNA, intact VLPs, or 20 ng of poly(I:C) HMW as a positive control (InvivoGen, #tlrl-pic-5). TLR3 activation induced by the RNA or VLPs was normalized to the response of poly(I:C)-stimulated cells.

### Statistical analysis

All data are presented as mean ± SEM. Data were analyzed using one-way ANOVA test or using Student’s *t*–test. Analyses were performed using GraphPad PRISM 10.6.1 (GraphPad Software Inc). The value of *p* < 0.05 was considered statistically significant. Statistical significance is noted in figures as \**p* < 0.05, \*\**p* < 0.01, \*\*\**p* < 0.001, \*\*\*\**p* < 0.0001.

## Data availability

All data are available from the corresponding author upon reasonable request. Source data are provided with this paper.

## Acknowledgements

This work was supported by grant from the Latvian Council of Science under Grant lzp-2022/1-0629.

## Contributions

**Anete Ogrina-Komarova:** Data curation; Formal analysis; Validation; Investigation; Methodology; Visualization; **Ieva Kalnciema:** Data curation; Formal analysis; Validation; Investigation; Methodology; **Dace Skrastina:** Resources; Data curation; Formal analysis; Validation; Investigation; Methodology; **Arnis Strods:** Data curation; Formal analysis; Validation; Investigation; Methodology; Conceptualization; Resources; Funding acquisition; Writing—original draft; Project administration; Writing—review and editing. **Santa Pikure:** Data curation; Investigation; Methodology; Visualization; **Juris Jansons:** Data curation; Investigation; Methodology; Visualization; **Gunta Resevica:** Data curation; Investigation; Methodology; **Janis Bogans:** Resources; Data curation; Investigation; Methodology; **Ramona Petrovska:** Resources; Data curation; Investigation; Methodology; **Andris Zeltins:** Resources; Formal analysis; Investigation; Methodology; **Ina Balke:** Conceptualization; Resources; Data curation; Formal analysis; Supervision; Funding acquisition; Validation; Visualization; Writing—original draft; Project administration; Writing—review and editing.

## Ethics declarations

The authors state they have no competing interests or disclosures.

